# Structural and evolutionary analysis unveil functional adaptations in the promiscuous behavior of serum albumins

**DOI:** 10.1101/2021.05.01.442285

**Authors:** Ana Julia Velez Rueda, Guillermo Ignacio Benítez, Leandro Matías Sommese, Sebastián M. Ardanaz, Estefanía L. Borucki, Nicolas Palopoli, Luis E Iglesias, Gustavo Parisi

## Abstract

Promiscuous activities have been related to the capacity to catalyze reactions different from those a protein has evolved to sustain. From this evolutionary perspective, we rethought the serum albumins promiscuous behavior. We found that the cross aldol condensation of acetone and p-formylbenzonitrile is a promiscuous reaction conserved in humans and other mammals. Structural and evolutionary analysis indicates that the involved residues could have evolved for a still unknown biological function. Our results could contribute to better characterize the serum albumin family and raise questions about the evolution of protein promiscuity and function.

## Introduction

First hypotheses on the origin of protein promiscuous activities proposed them as the starting point for new protein activities ^1^. Many directed evolution studies showed that protein plasticity ^2^, a feature of many promiscuous proteins and complexes ^3^, is an important factor for random mutations to endure selection throughout generations. In the same sense, protein backbone rearrangements could be responsible for changes in the protein's functionality ^4^. These observations may be expected taking into consideration the relationship between conformational diversity, biological function and evolvability ^5^.

Tawfik and co-workers defined promiscuity as the capability of a protein to catalyze reactions different from those it has evolved to sustain ^6^. From this evolutionary perspective many cases of promiscuous behavior in proteins, and their functionality in general, could be rethought. The serum albumins constitute one of these very interesting cases of catalytic promiscuity that need to be revisited. Members of this family show a large structural diversity in spite of global sequence conservation ^7^. Moreover, they differ in the ability to perform complex promiscuous activities, their catalytic ability has been tested for several reactions ^8–12^ and they have also been used as biocatalysts in organic reactions such as the Henry reaction, the Kemp elimination and the cross aldol condensation, to give a few examples ^10,13,14^.

Human Serum Albumin (HSA) is the main protein in plasma, binds multiple ligands ^15^, and has recently emerged as a very important drug carrier ^16,17^. This single-chain protein has several high-affinity binding sites, however the majority of the drugs and ligands bind to the so named Sites I and II ^18^. In particular, residues Lys 199, Arg 410, Tyr 411, Cys 34, and Lys 195 from HSA are described as some of the important ones, not only for ligand binding but also for catalysis ^19–21^. Among the best characterized catalytic activities described are the esterase-like activities and the thioesterase activity, shared with other albumins as Bovine Serum Albumin (BSA) ^9^. Such multiple catalytic capacities can be associated with some structural cavity features such as the presence of activated amino acids (with abnormal pKas) in mostly hydrophobic environments, which would increase the catalytic repertoire favoring the attack of substrates and allow creating the microenvironment suitable for catalysis ^22,23^. The structural determinants that allow such a diversity of reactions to be carried out in the same active site of albumins, with minimal variations between species, are still as poorly understood as the biological implications and evolutionary origin of such activities. If promiscuous activities arise spontaneously and due to structural or mechanistic similarity, there should be no evolutionary imprint that demonstrates functional adaptation^2,24^.

Given this and due to the numerous promiscuous reactions described for albumin, we wonder if these activities occur by chance or if they actually have physiological or adaptive bases that remain as yet uncharacterized. Is albumin, therefore, an enzyme of activity not yet characterized? In the present work, we analyze structural and evolutionary patterns of HSA, BSA, RabSA (Rabbit Serum Albumin), PSA (Pig Serum Albumin), and RSA (Rat Serum Albumin) as representatives of the albumin family to bring some light to these questions. For these purposes, we used evolutionary and structural analysis to characterize the probable presence of functional adaptations during evolution in positions previously reported as supporting the promiscuous behavior in albumins. To evaluate this hypothesis we used the cross aldol condensation of acetone and *p*-formylbenzonitrile, previously assayed with BSA ^14^, now extended to other members of the albumin family.

## Material and Methods

### Evolutionary Analysis

Aiming to analyze if the promiscuous activities of albumins are physiologically relevant reaction mechanisms, we decided to study the evolutionary bases of the promiscuous reaction aldol condensation, characterized in our previous works ^12,14^. The coding gene sequences and homologous proteins used for this purpose were obtained using BLAST ^25^, using human serum albumin like query. and the multiple sequence alignment of serum albumins was constructed using the Clustal algorithm ^26^. A total number of 46 representative sequences of different organisms, with known structures (121 total structures in the PDB ^27^), were included in the analysis (see Supplementary Table 1). The evolutionary tree was estimated using Phyml ^28^ with the JTT model of evolution, experimental frequencies, and a gamma distribution of rate variation among sites. Also, nonparametric bootstrapping with 100 replicas was applied. To estimate the sites subject to positive selection, we used the mixed-effects model of evolution (MEME) ^29^ available for execution in the Datamonkey service ^30^. The residues under positive selection were mapped to the structures in order to estimate structural and functional relationships.

### Ancestral Sequence Reconstruction

Ancestral sequences were reconstructed using PAML version 4.9 and the Lazarus package ^31^, incorporating the gamma distribution of rate variation and JTT as substitution model. For each site of the inferred sequences, posterior probabilities were calculated for all 20 amino acids and the residue with the highest posterior probability was then assigned at each site.

### Evolutionary History Simulation

A simulation of the evolutionary history was done using the Seq-gen software ^32^. We used the same tree obtained by phylogenetic inference (see ‘Evolutionary Analysis’) with the JTT evolutionary model and a sequence rich in “KK” pairs as sequence seed, in order to calculate the random frequencies of occurrence of “KK”, “KR”, “RK” and “RR” pairs in the tips of the phylogenetic tree.

### Structural Analysis

We used Fpocket ^33^ to identify the cavities in the available conformers (see Supplementary Table 2) with known structures of all proteins in the dataset. We estimated the pKa values of the ionizable residues of these conformers using PROPKA ^34^, to identify relevant amino acids for biological function or promiscuous activity. The residues predicted as being under positive selection were mapped to the cavities and assigned a pKa when possible. Preparation and analysis of data were done using custom scripts in the Python programming language.

### Albumins cross aldol condensation activity test

HSA, RabSA, RSA, PSA and BSA were tested for the cross aldol condensation between acetone and *p*-formylbenzaldehyde (using the experimental conditions) following the protocol setted up for bovine serum albumin (BSA) catalysis in previous work ^14^. According to this protocol, a mixture of 54 mg of the assayed albumin was added to a solution of *p*-formylbenzaldehyde (0.05 mmoles) in acetone-ethanol 1:1 (1 ml). The resulting mixture was shaken at 45 °C and 200 rpm; at different times, aliquots from the biotransformation were taken and after centrifugation of the albumin, submitted to HPLC analysis as reported (10). The conversions to enone (cross aldol condensation product) obtained from each albumin and determined by HPLC are shown in Figure 1. No appreciable conversion was detected in absence of protein, as already observed by us ^14^.

**Figure 1.**
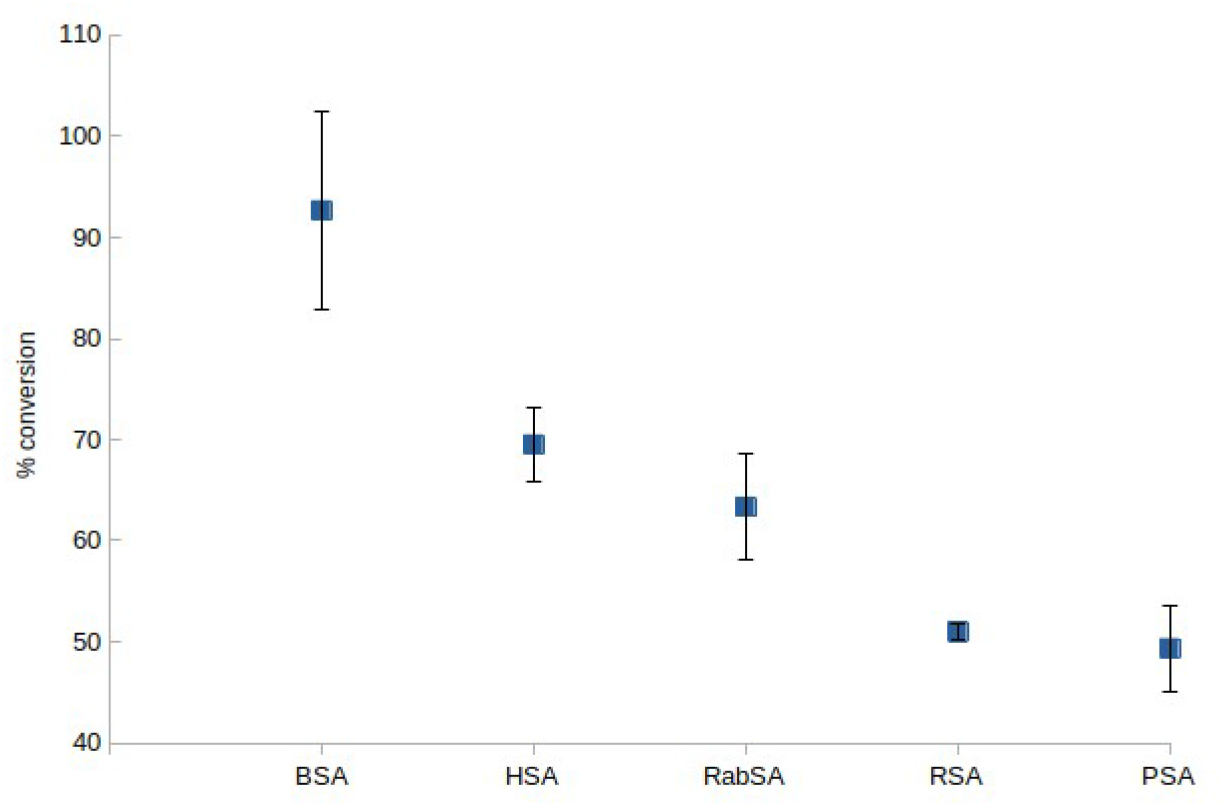
Conversion to enone (cross aldol condensation product) in the reaction of acetone with *p*-formylbenzonitrile catalyzed by albumin from different species. The biotransformations and their analysis were conducted following reported protocols 14. Conversion is represented qualitatively in the figure by a cross, being more significant in BSA and the lowest activity being observed in RSA and PSA serum albumins.

## Results

### Conserved promiscuous behavior of albumins

We have recently characterized the capacity of BSA to sustain the promiscuous cross aldol reaction involving a set of ketones and benzaldehydes ^12,14^. Here, we have extended the study of the reaction of acetone and p-formylbenzaldehyde to human (HSA), rat (RSA), rabbit (RabSA), and pig (PSA) serum albumins. Besides the sequence divergence of the site involved in such reaction, these five albumins were able to catalyze the reaction (Figure 1). In Figure 2 we represent a sequence alignment of residues involved in the cross aldol condensation following previous characterization ^14^. Following HSA numeration, Lys 199, Arg 218, and Arg 222 are residues involved in this reaction located in the second largest cavity detected by Fpocket in the domain AII as we characterized in previous works ^14^. These residues are highly conserved as we can observe in the alignment of Figure 2.

**Figure 2.**
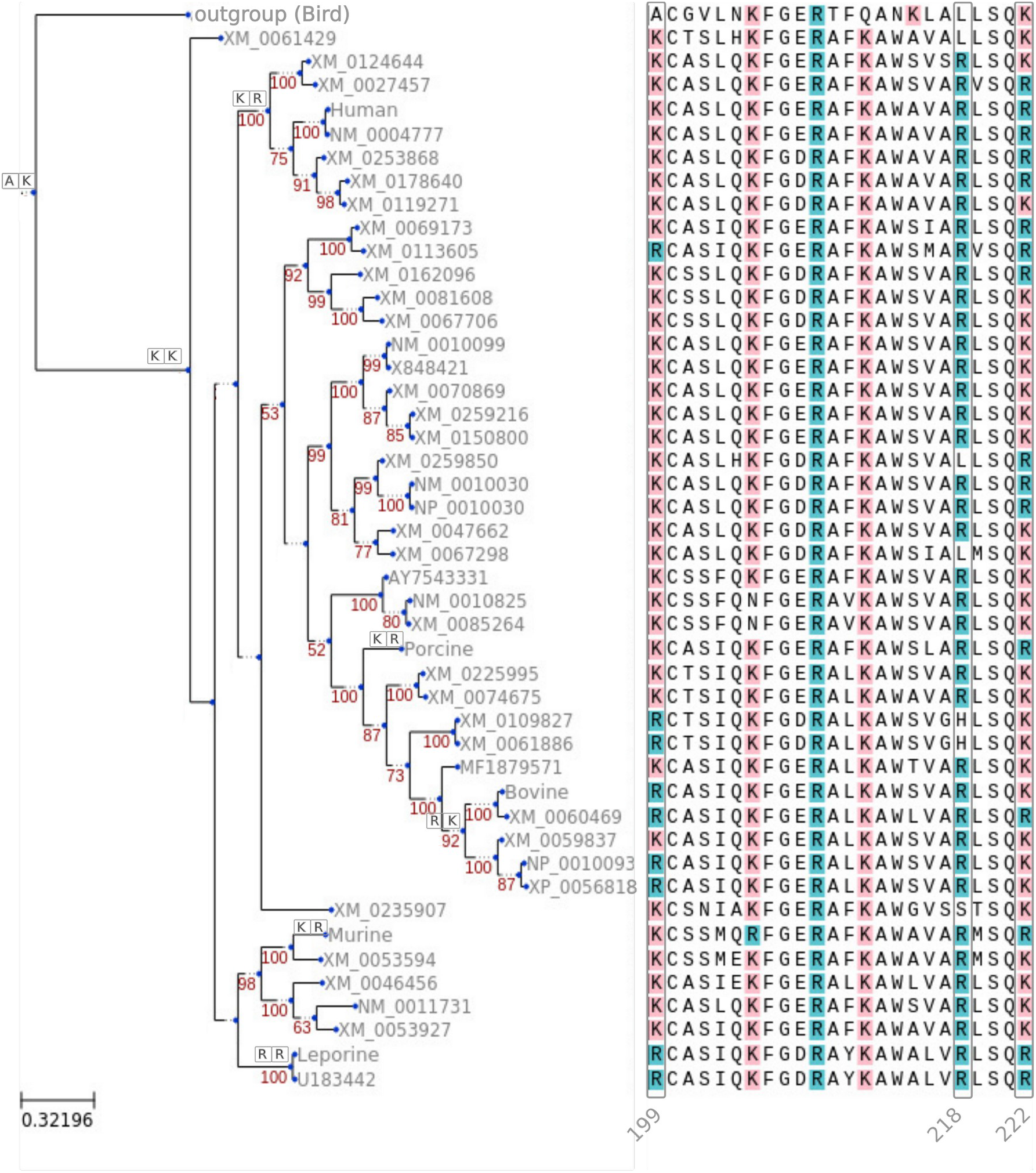
Multiple sequence alignment of serum albumin proteins from 46 different species (including pigeon seroalbumin, used to root the tree) showing relevant portions of the HSA sequence involved in the aldol condensation. Arginine and lysine are colored blue and pink respectively. Also, the residues involved in the aldol condensation, Lys 199, Arg 218, and Arg 222 (following the HSA numeration) were highlighted. In the internal nodes are highlighted the evolutionary changes (derived from the ancestral reconstruction protocol, see Methods) in positions 199 and 222, respectively. No changes were observed on position 218..

As derived from Figure 2, it is interesting to note the shift between Lys 199 and Arg 222 in HSA to the equivalent residues observed in BSA (Arg 198 and Lys 221 according to BSA numeration). Apparently, those replacements are conservative since both residues could be positively charged at neutral pH. However, as will be described below, their pKas are very different and could affect their role in catalyzing the reaction. Also, in PSA, both equivalent positions are occupied by Arg. Besides these replacements, the five seroalbumins showed catalytic activity for the tested reaction (see Figure 1), confirming the conservation of amino acid replacements.

### Positive selection analysis and cavities

As a first step in the sequence analysis, we estimated the presence of protein residues evolving under a positive selection process. These residues are thought to be involved in functional adaptation ^35,36^. To achieve this goal, we built a multiple sequence alignment of 46 albumin proteins from 38 different species (see Supplementary Table 1). In particular, 6 representatives have known crystallographic structures. Using the MEME algorithm we detected a total number of 38 sites evolving under positive selection on the alignment (Supplementary Table 2).

Aiming to have a deeper understanding of the biological relevance of these positions, we mapped these residues to the cavities predicted by Fpocket in all the conformers of the different proteins with known structures. We also complemented the biological data using the information about functionally important residues and cavities, collected from the bibliography ^17,37–39^. From this mapping we observed the overlap of approximately 50 to 70% of positively selected residues with the biological relevant cavities (Figure 3 and Supplementary Table 2), fluctuating according to the species and conformations used. Among these cavities we found the IIA drug binding site, described previously ^40^ as containing residues involved in promiscuous reactions in HSA and BSA. This cavity is one of the major cavities predicted by Fpocket and contains the residues involved in supporting the aldol condensation ^41^. Interestingly, Arg 222 is evolving under a positive selection process, suggesting a putative functional role (Supplementary Table 2).

**Figure 3.**
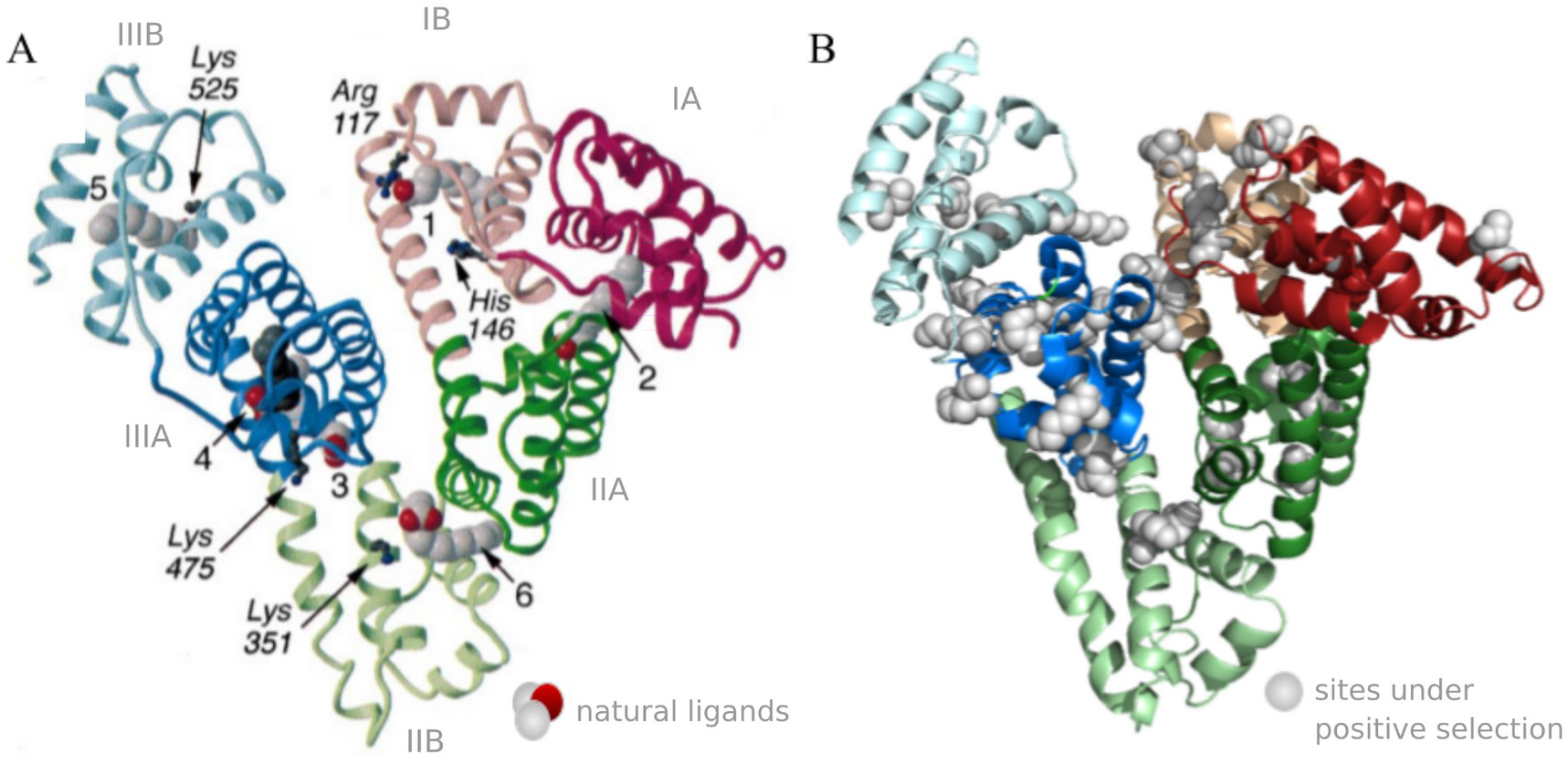
A) Natural ligand binding sites on human serum albumin (HSA). The serum albumins domains I to III and their subdomains (A and B) are differentially colored and depicted. Additionally, the approximate locations of the natural ligand binding sites of human albumin are highlighted. Figure adapted from 42. B) Sites under positive selection predicted with MEME software, exemplified on the crystallographic structure of human albumin (PDB ID 1AO6).

The high percentage of overlapping residues evolving under positive selection and biological relevant cavities indicate that most of them are participating in the transport, binding and/or catalytic promiscuous activities, indicating a recent pattern of functional adaptation in the evolution of seroalbumin.

### Analysis of pKa shifts

It is well established that pKa values of residues can shift their values due to different structural and environmental features ^43^. Residues with pKa shifts could be involved in critical roles in catalytic mechanisms, protein-protein interactions, and ligand binding and could be associated with the support of several biological functions ^44^.

We analyzed the occurrence of pKa shifts as a proxy to detect residues involved in the biological process. From the analysis of the pKa values of the ionizable residues in all the available albumins conformers with known structure, we were able to detect the presence of activated amino acids in the cavity where the putative active site occurs in HSA. In particular, residues Lys 199 and Arg 222 described before, showed an abnormal acid pKa (see Table 1).

**Table 1.** Summary table of pKa shifts in positions 199 and 222 presumably involved in catalysis, specified by species (HSA, BSA, and RabSA)

| Protein | Residue, position, observed pKa shift |  |
| --- | --- | --- |
| HSA | Lys, 199, ~7.51 | Arg, 222, ~9.49 |
| BSA | Arg, 199, ~9.24 | Lys, 221, ~7.27 |
| RabSA | Arg, 199, ~16.53 | Arg, 222, ~8.25 |

This activation seems to be related to the presence of basic residues that are in contact with the mentioned residues. We observed that position Lys 199 of HSA is abnormally acidic (average pKa among the conformers is ~7.51, see Supplementary Table 2 information about conformers used) while Arg 222 has a pKa ~9.49, also abnormally acid for an Arg residue. When we analyzed equivalent positions in BSA, we found that Arg 198 has a pKa ~9.24 (equivalent to Lys 199 of HSA). However, Lys 221 (equivalent to Arg 222 of HSA) shows a pKa ~7.27. As reported previously ^12,14^, an acidic residue acting as a nucleophile is required to sustain the aldol condensation of acetone with p-formylbenzonitrile. This acid residue is postulated to be the Lys 199 in HSA and Lys 221 in BSA. In this sense, the “active site” is inverted in these two seroalbumins. As we discussed early, these positions are well conserved due to the observed substitutions of Lys and Arg, residues with positive charges that can be normally interchanged during evolution. However, physicochemical constraints related to the containing cavity introduced pKa shifts, although inverted, to support the promiscuous reaction. More interestingly, equivalent positions to Lys 199 and Arg 222 in HSA are occupied by two Arg in RabSA. The analysis of their pKa indicated that the amino-terminal Arg (Arg 199) showed an abnormal basic shift with a pKa ~16 while the carboxy-terminal Arg (Arg 222) has a pKa ~8. However, according to the experimental results shown in Figure 1, an acid Arg and a basic Arg sustain the reaction with much less efficiency than HSA and BSA, a fact which could explain taking into account the lesser nucleophilicity of arginine compared with lysine. Unfortunately, the other serum albumins used, PSA and RSA, lack a known crystallographic structure and pka shift estimation is not possible.

### Phylogenetic Analysis

The inversion of the “active site” configuration between HSA and BSA mentioned before is evidence of the evolutionary conservation of acid-basic properties required to sustain the catalytic condensation of the aldol. How many times did this inversion occur in the evolutionary history of the seroalbumin studied? To answer this question we used the phylogenetic tree in Figure 2 and applied ancestral reconstruction techniques to infer ancestral states for the equivalent positions to Lys 199 and Arg 222 from HSA (represented as “KR”). As it is derived from Figure 2, we show in given internal nodes the estimated ancestral states for those positions. For example, the ancestral node involving all mammals contains two Lys in equivalent positions to Lys 199 and Arg 222 (shown as “KK” in the tree of Figure 2). Among the organisms in our phylogenetic tree, we can find different combinations like “KR” (*Homo sapiens, Sus scrofa, and Rattus norvegicus*), “RK” (*Bos bovis and Capra hircus*), or “RR” (*Chinchilla Lanigera, Oryctolagus cuniculus, and Bubalus bubalis*). The occurrence of these shifts of amino acid with the conservation of the corresponding pKa shifts possibly indicates the biological importance of these residues.

Finally, for a deeper analysis, we estimated the random occurrence of these shifts using the phylogenetic tree in Figure 2 and setting the ancestral state of these two positions as “KK” as derived from the ancestral reconstruction results. Then we simulated the evolutionary history of sequences using Seq-gen ^32^ and counted the occurrence at the tips of the different combinations. Comparing the frequencies in the natural sequences and in those simulated with Seq-gen we obtained that the occurrence for KR and RK pairs are significantly greater (p-value < 0.001) in the observed sequences than in the simulated sequences. Our result suggests that this inversion is structurally constrained indicating a putative functional adaptation.

### Tunnels conservation

In the last years, attention to the presence of tunnels has increased due to their importance to explain structure-function relationships in proteins ^45^. Tunnels allow the transit of ligands from the outside (and vice versa) of the protein to inner cavities containing active and binding sites. Overall, their structure and physicochemical properties define binding constants for ligands. Along with protein dynamism, tunnels also define active and inactive conformations just allowing the opening or closing of cavity access ^45–47^. Using the program MOLE, we identified a common tunnel linking Lys 199 and Arg 222 with the surface in HSA and their structurally equivalent residues in BSA and RabSA. At the bottom of the tunnel, we also found the conserved residue Arg 218 which is spatially close to Lys 199 and Arg 222 (Figure 4). Interestingly, at the entrance of the tunnel, we found a mostly conserved positively charged residue (His, Arg, or Lys). This residue, an His in HSA (His 440), along with Arg 218 are evolving under a positive selection process involved in functional adaptations. Our hypothesis is that this tunnel common to HSA, BSA, and RSA allows the transit of biologically important ligands up to the cavity where Lys 199 and Arg 222 are found.

**Figure 4:**
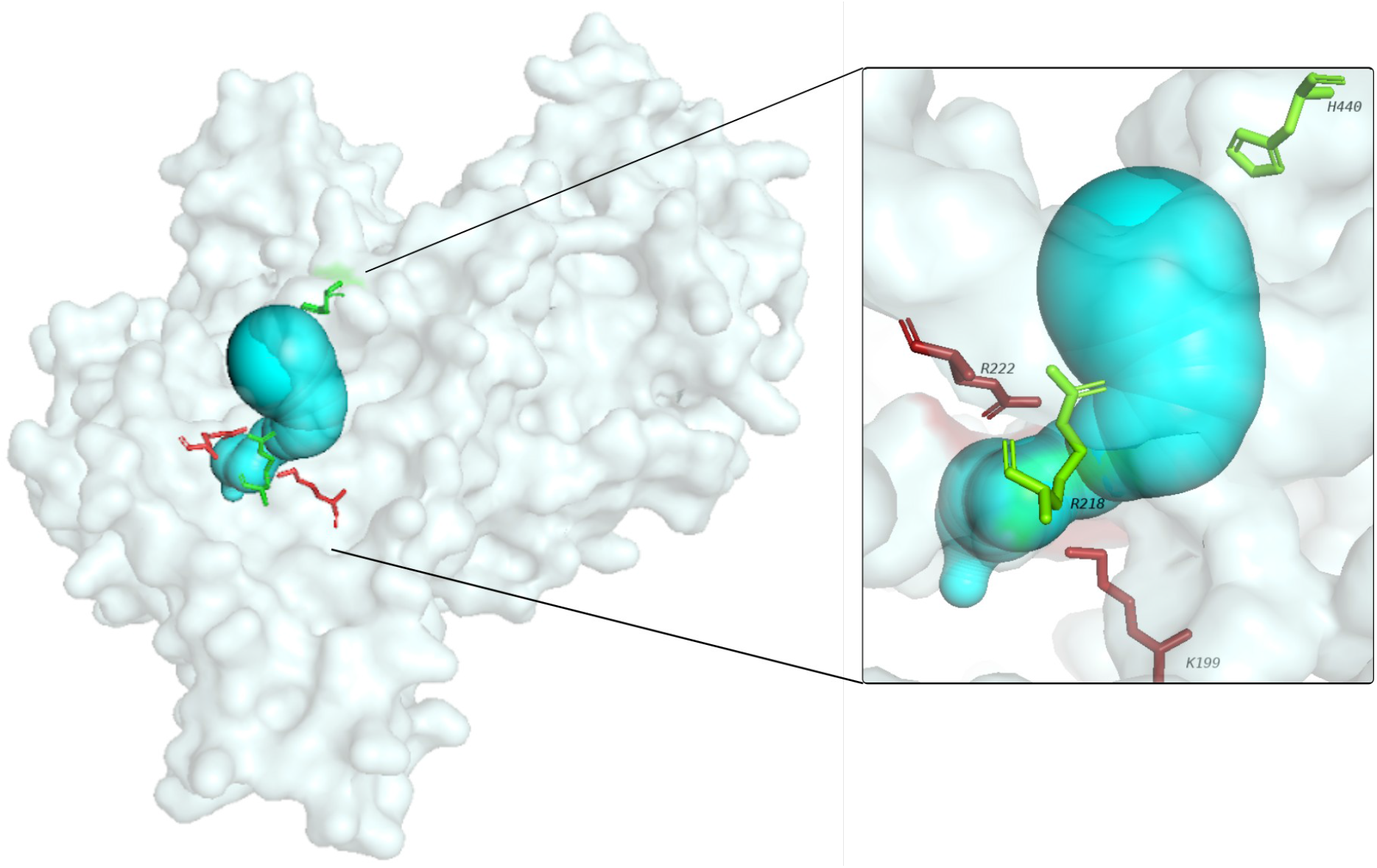
Surface representation of HSA (PDB 1ao6) showing the tunnel connecting the surface of the protein with the residues involved in the aldol condensation. In red, in stick representation, Lys 199 and Arg 222. In green, Arg 218 in the same cavity and spatially close to the residues mentioned before. In the entrance of the tunnel, we found a conserved positively charged residue (His, Lys, Arg) evolving under positive selection.

## Discussion

In summary, we have collected different pieces of evidence that support the view that the residues required to sustain a promiscuous reaction in serum albumin could have evolved to hold a biological function. We have found that those residues involved in the reaction (Lys 199 and Arg 222 in HSA) are located in a large and hydrophobic cavity and are conserved during evolution (Figure 2) Interestingly, the pka shifts in these amino acids are also conserved to produce polarized “acid” and “basic” residues independently of being Lys or Arg. As we previously showed, the swapping in the Lys/Arg (HSA) and Arg/Lys (BSA) key residues (Table 1) in the natural sequences analyzed is statistically different from the occurrence by chance of the simple replacements of Lys and Arg. Evolutionary models as those used in this work (see Methods) consider residue replacements during evolution in a site-independent fashion. However, conservation of pka shifts in species also indicates physicochemical environmental constraints indicating a putative functional adaptation.

Another piece of evidence is the finding of a common tunnel in the three structures analyzed. Tunnels allow the transit of ligands between cavities and protein surfaces. As shown in Figure 4, three important residues lie at the bottom of the tunnel (Lys 199, Arg 218, and Arg 222 in HSA numbering). Among these, Lys 199 and Arg 222 were previously suggested to be involved in promiscuous reactions ^41,48,49^. Also, Arg 218 is 100% conserved in serum albumins and its replacement has been related to the occurrence of human diseases. Replacement of Arg 218 to His or Pro produces abnormal albumin with increased affinity for serum thyroxine found in an autosomal dominant condition called Familial dysalbuminemic hyperthyroxinemia ^50,51^. This condition is caused by an abnormal albumin molecule with an increased affinity for the hormone thyroxine. Supporting the putative role of this tunnel is the fact that two flanking residues (Arg 222 and His 440 in HSA see Figure 4) are evolving under positive selection.

Our results offer a biological interpretation of the observed promiscuous catalytic activity in serum albumins. In addition, a new perspective is offered in the classification of albumins as possible enzymes, in light of the observation of the selective pressure in the key positions for catalysis, typically observed during functional divergence ^52^. From this point of view, the catalytic properties of the serum albumins documented in the present work could be involved in an unknown biological process rather than a promiscuous behavior. These results not only allow us to better characterize the family but also open up interesting questions about the origin of promiscuous behavior and the evolution of protein function.

## Acknowledgments

GP, LMS, LEI, and NP are researchers, and AJVR and GIB are Ph.D. fellows from CONICET. This work was supported by Universidad Nacional de Quilmes (PUNQ 1004/11), ANPCyT (PICT-2014-3430, PICT-2013-0232). The funders had no role in study design, data collection, and analysis, decision to publish, or preparation of the manuscript.

## Authors contributions

AJVR and GP conceived the study and were in charge of overall direction and planning. AJVR, GIB, and GP carried out the evolutionary and structural analyses. LMS carried out the statistical analyses and plottings. SA and ELB performed the experimental tests. AJVR and GP wrote the manuscript with input from all authors. LEI and NP collaborated with the revision of the manuscript.

## Supplementary Tables

**Supplementary Table 1.**
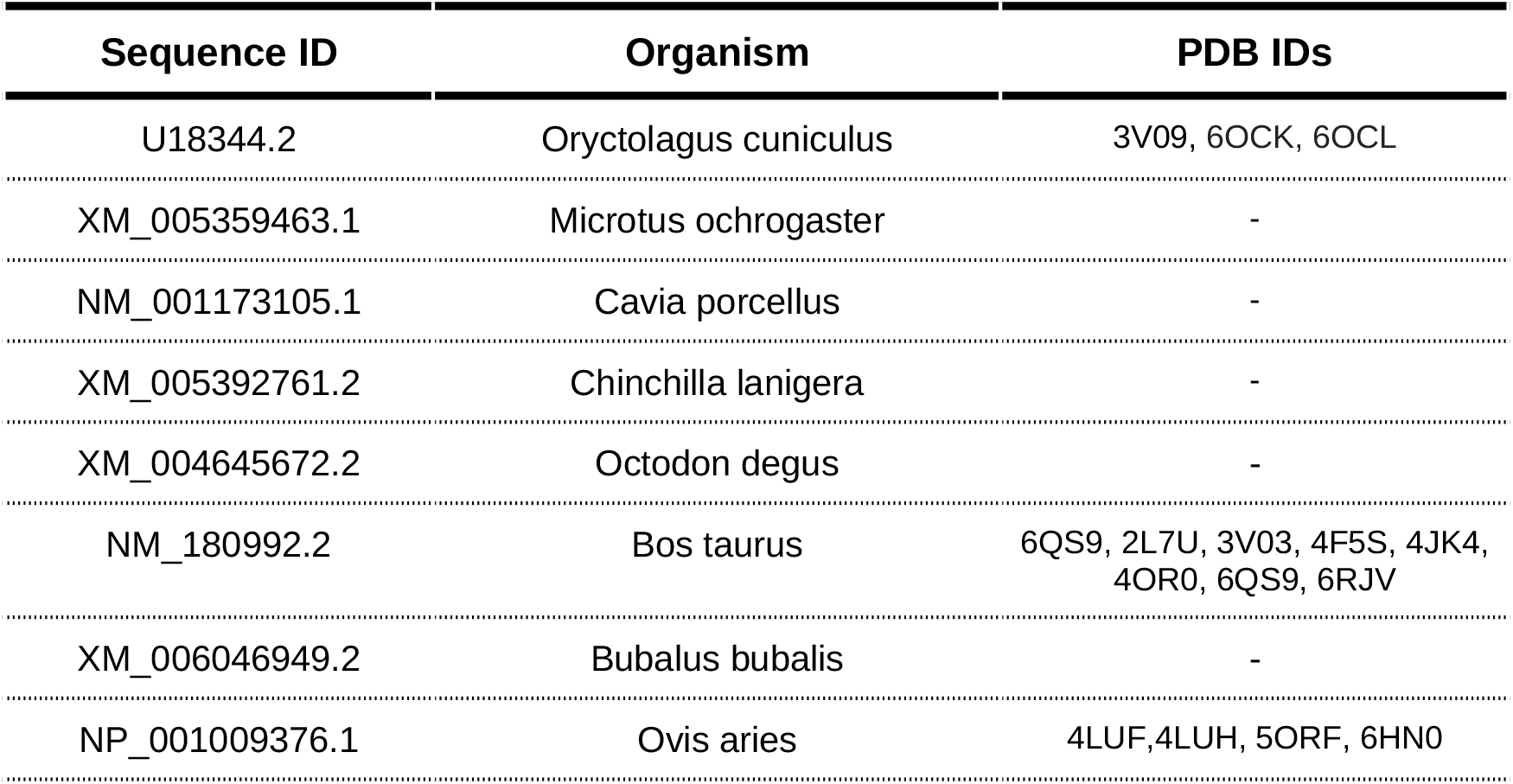
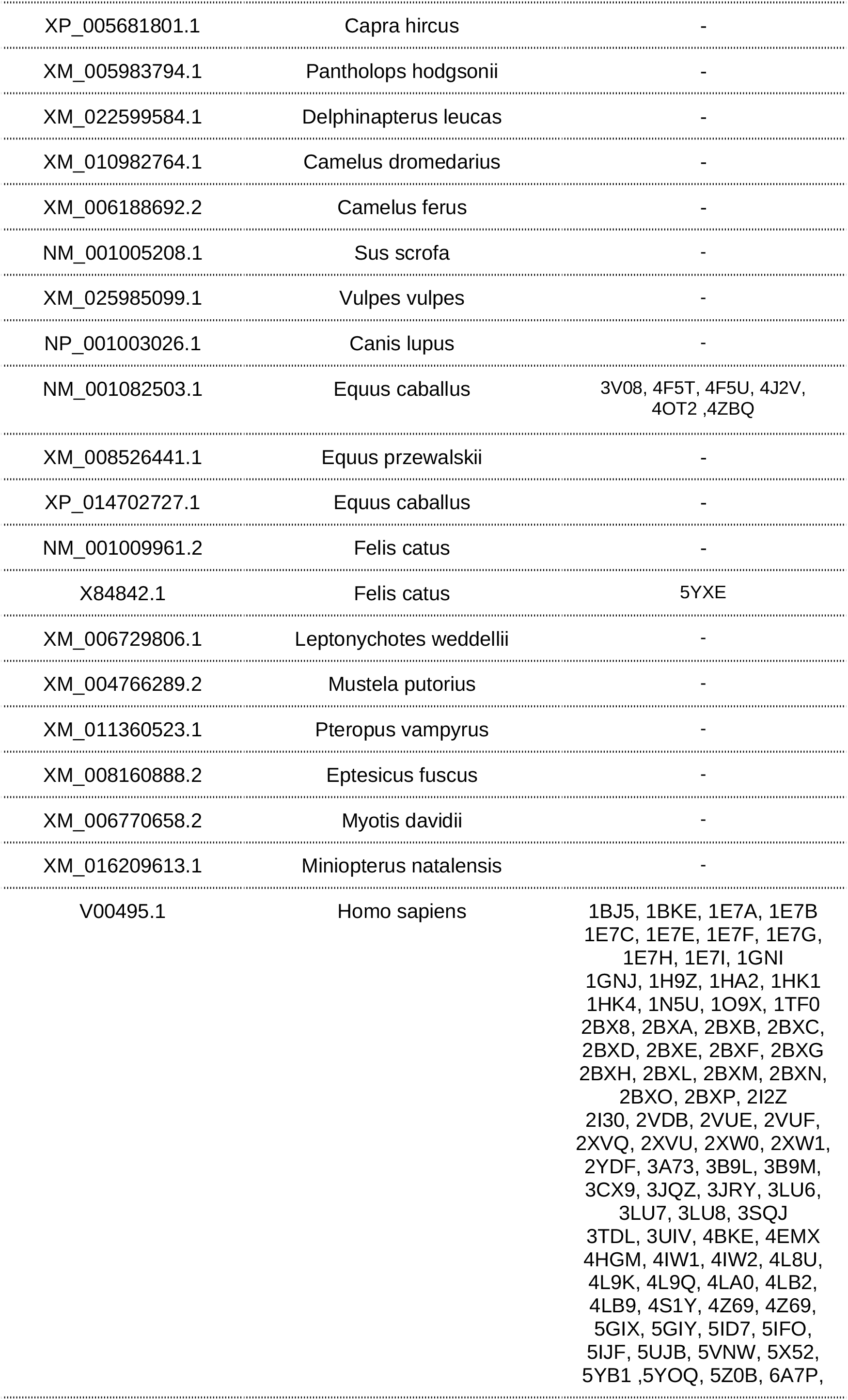
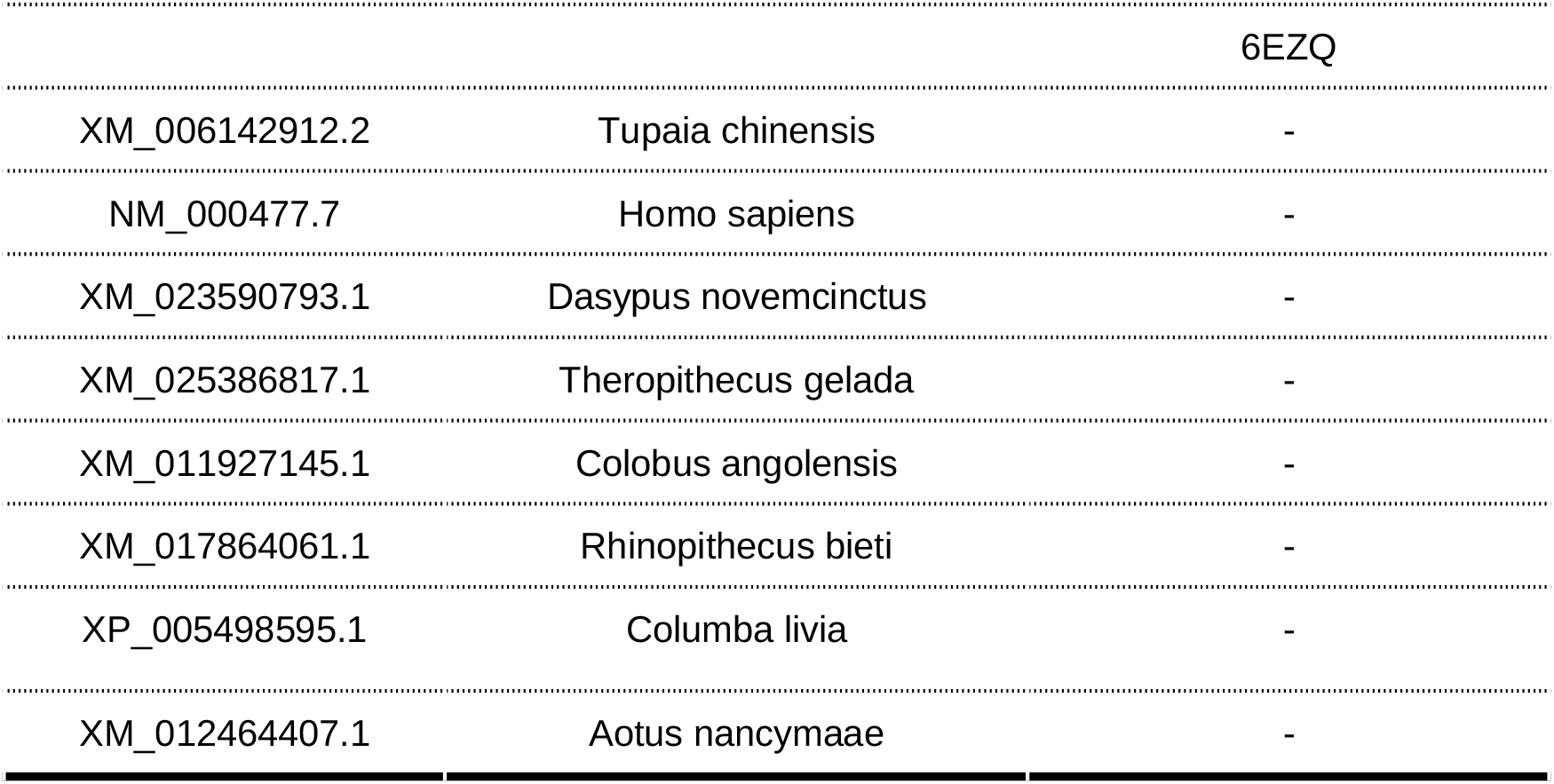
Alignment sequences and their corresponding information about structures and organisms

**Supplementary Table 2.**
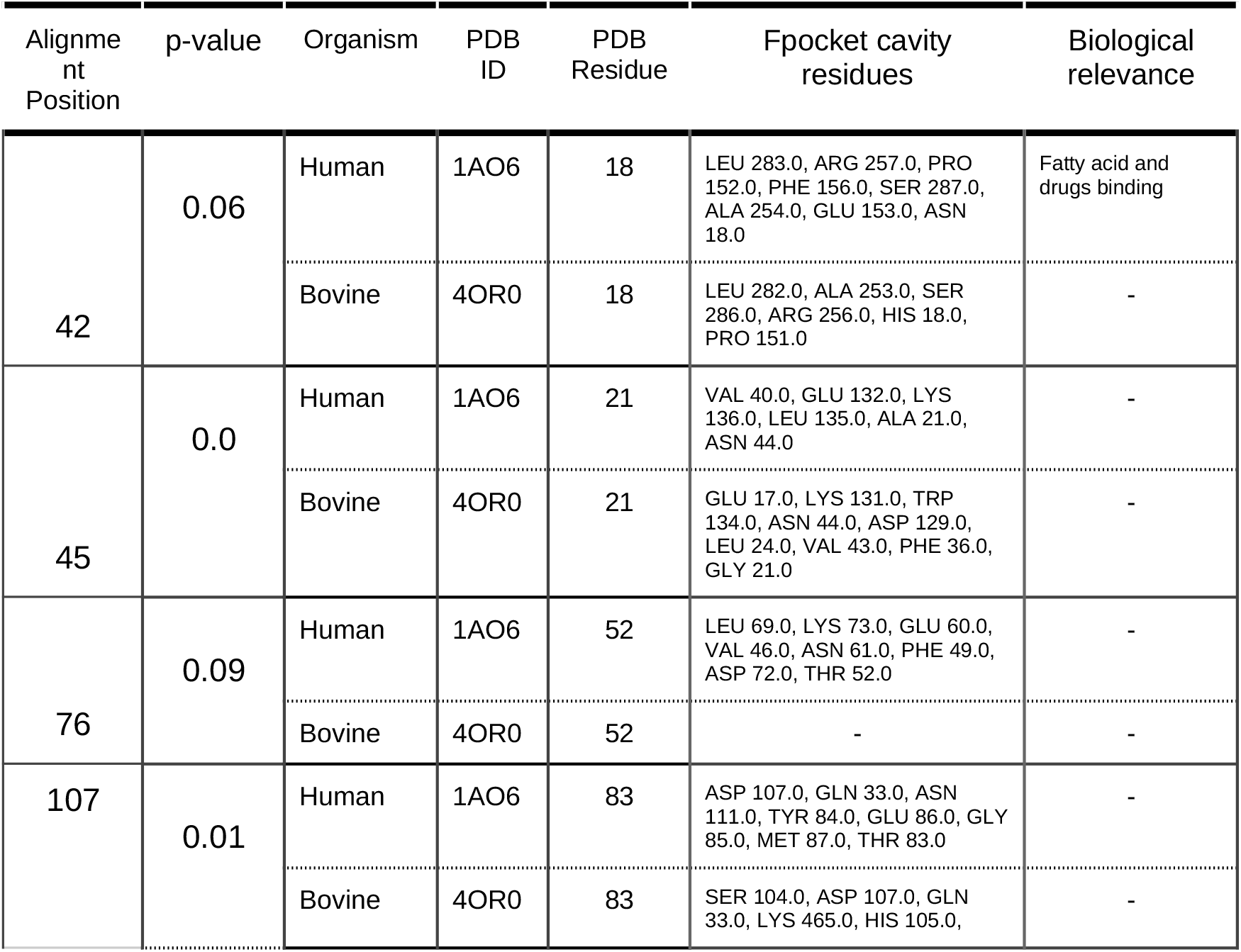
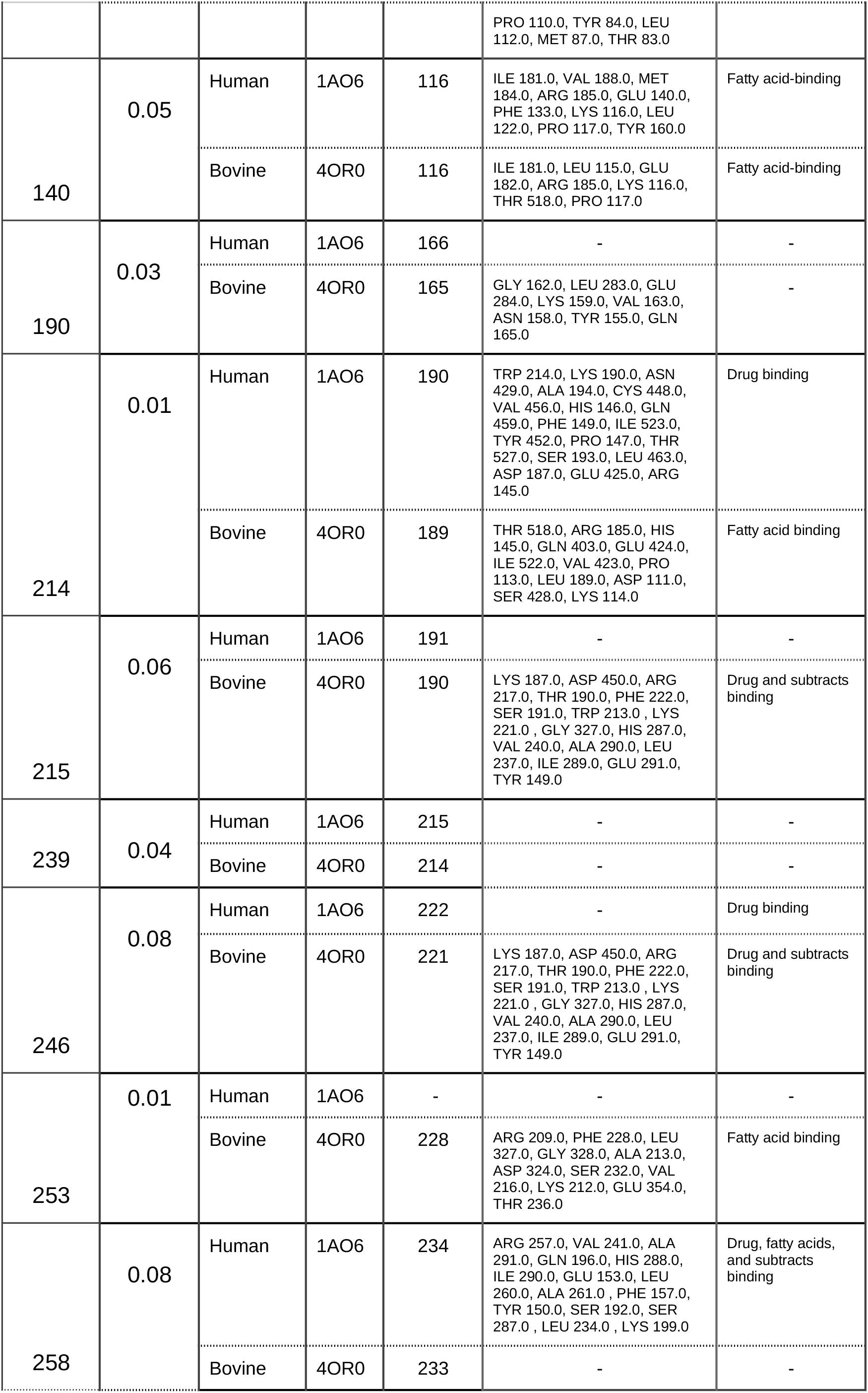
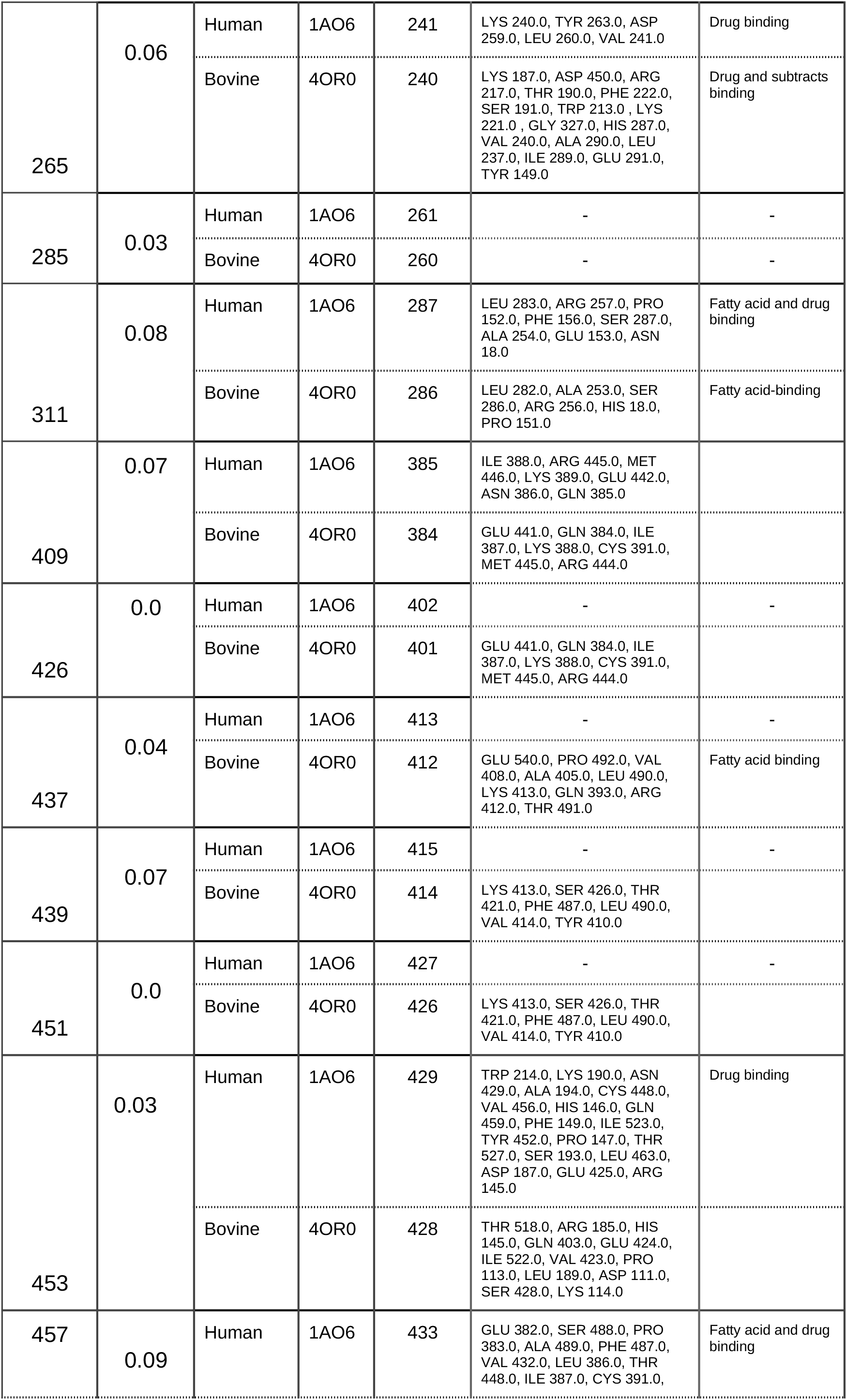
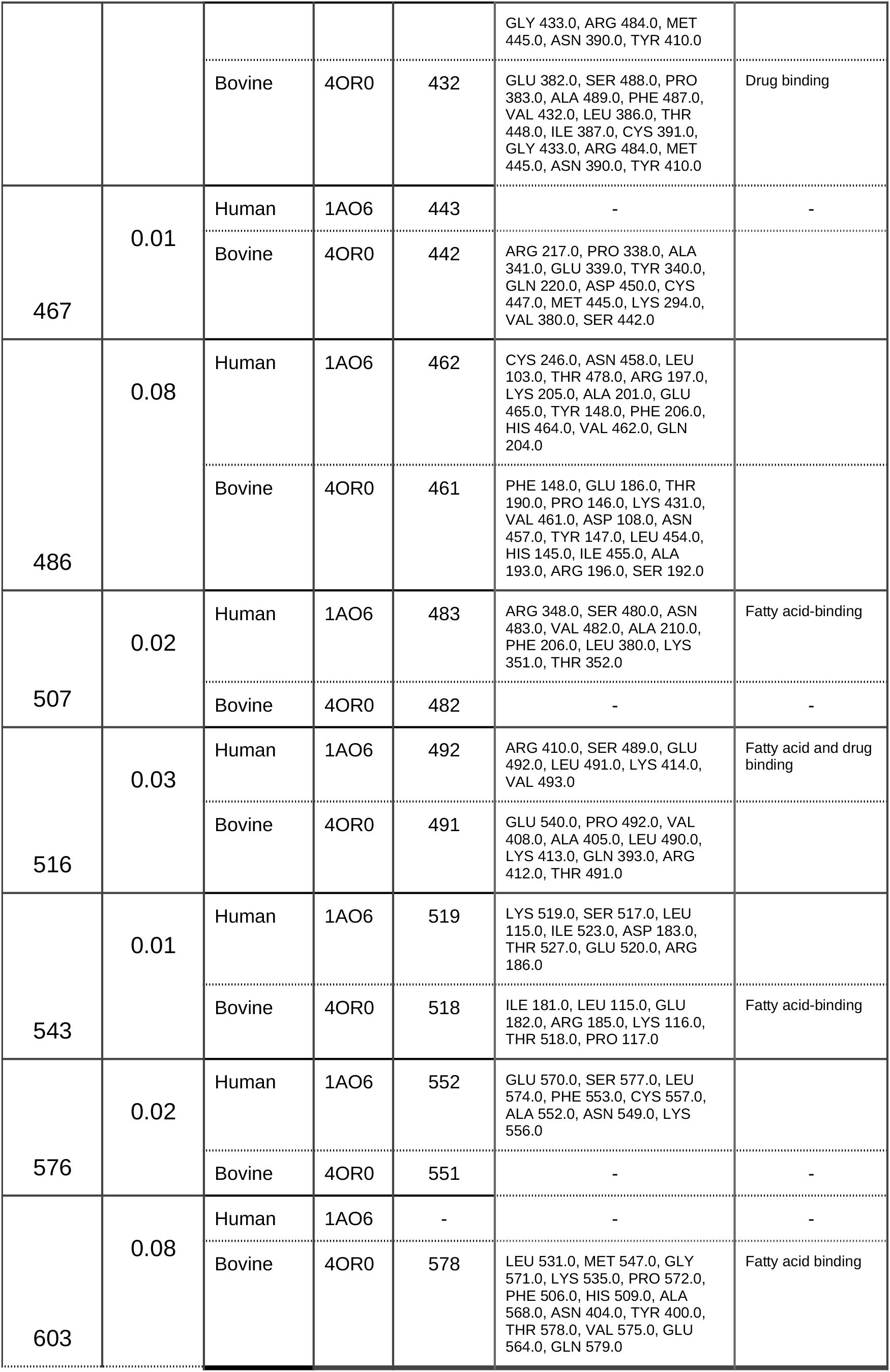
Positively selected residues referenced to a Bovine and Human crystal structure. The selected positions detected by MEME ^29^ in the alignment, and their related positions in the protein PDB structures are detailed in the table, as well as the biological relevance of each residue and cavity. Positions in the alignment which showed positive selection but have no structural information are not included in the table. Biological relevance was collected from previous works ^16,19,38,41,53–58^.

